# CX3CL1 binding protein-2 (CBP2) of *Plasmodium falciparum* binds nucleic acids

**DOI:** 10.1101/662494

**Authors:** Ritu Saxena, Jasweer Kaur, Rachna Hora, Palwinder Singh, Vineeta Singh, Prakash Chandra Mishra

## Abstract

Several exported *Plasmodium falciparum*(Pf) proteins contribute to malaria biology through their involvement in cytoadherence, immune evasion and host cell remodelling. Many of these exported proteins and other host molecules are present in iRBC (infected red blood cell) generated extracellular vesicles (EVs), which are responsible for host cell modification and parasite development. CX3CL1 binding proteins (CBPs) present on the surface of iRBC have been reported to contribute to cytoadhesion by binding with the chemokine ‘CX3CL1’ via their extracellular domains. Here, we have characterized the cytoplasmic domain of CBP2to understand its function in parasite biology using biochemical and biophysical methods. Recombinant cytoplasmic CBP2 (rcCBP2) binds nucleic acids showing interaction with DNA/RNA. rcCBP2 shows dimer formation under non-reducing conditions highlighting the role of disulphide bonds in oligomerization while ATP binding leads to structural changes in the protein. *In vitro* interaction studies depict its binding with a Maurer’s cleft resident protein ‘PfSBP1’, which is influenced by ATP binding of rcCBP2. Our results suggest CBP2 as a two-transmembrane (2TM) receptor responsible for targeting EVs and delivering cargo to host endothelial cells. We propose CBP2 as an important molecule having roles in cytoadherence and immune modulation through its extracellular and cytoplasmic domains respectively.

## 1. Introduction

*Plasmodium falciparum* (Pf) replicates asexually within iRBCs to manifest the symptoms of malaria [1]. During this stage of the parasite life cycle, Pf exports several proteins including families like PfEMP1, Rifin, Stevor and Surfin etc. to the host cell that are collectively termed as the malaria exportome [2]. These exported proteins remodel iRBCs to alter their rigidity and adhesive properties, facilitate nutrient uptake and help them to evade the host immune response [3]. Some exported proteins mediate binding of parasitized RBCs (pRBCs) with microvascular host endothelial cells via a phenomenon called as ‘cytoadherence’, with PfEMP1 being a key effector of the process [4]. Members of the PfEMP1 protein family are known to bind various host endothelial receptors like CD36, ICAM-1, VCAM etc. to cause sequestration in the microvasculature [5].

In 2003, Hatabu *et al* reported the chemokine CX3CL1 as another host endothelial membrane receptor for cytoadhesion of pRBCs in the brain of cerebral malaria patients [6]. CX3CL1/ fractalkine (FKN) expresses constitutively in a variety of non-hematopoietic tissues including brain, heart, lungs and kidneys, and exists both in membrane bound and soluble isoforms [7-8]. While its membrane bound form functions as an adhesion molecule, the soluble form exhibits chemotactic activity towards natural killer (NK) cells and monocytes [9-10]. Sargeant *et al* had identified a novel protein family termed ‘hypothetical 8’ with two iRBC surface expressed members [11] that were shown to act as parasite ligands involved in CX3CL1 mediated cytoadherence of iRBCs [12]. These were named CX3CL1 binding proteins 1 (CBP1; PF3D7_0113900) and 2 (CBP2; PF3D7_1301700) that share 32% sequence identity. Also, Hossain *et al* had predicted CBP2 to harbour a cold shock DNA binding domain and showed its nuclear localization in asexual parasites [13]. Also, localization studies on transgenics of CBP2 (Hyp8-GFP/ Hyp8-HA) have previously shown PEXEL (*Plasmodium* export element) motif containing CBP2 to co-localize with PfSBP1 at Maurer’s clefts (MCs) during the ring stage of parasite development [14]. Maurer’s clefts are parasite derived membranous structures that first appear in the iRBC cytosol in early ring stage of the parasitic development [15-16]. PfSBP1 and several other proteins including MAHRP1, Pf322 and REX1 & 2, are believed to be MC residents with roles in PfEMP1 transport [17]. Multiple deletion attempts on CBP2 gene in parasites were unsuccessful suggesting that it may be an essential protein [14, 18]. However, a recent study by Zhang *et al* using the *piggyBac* transposon approach showed that CBP2 is mutable [19]. Both PfSBP1 and CBP2 have also been reported to be present in extracellular vesicles (EVs) that are released by parasitized RBCs and have a significant relation with development of the disease [20-21]. These EVs carry a multitude of proteins and nucleic acids that are transferred to a variety of target cells including NK cells, macrophages and endothelial cells [22]. Pf derived EVs are involved in inter-parasite communications, development of gametocytes and regulation of host immune response [23-24].

In this study, we have cloned and purified the cytoplasmic domain of CBP2 to understand its role in iRBCs and Pf derived EVs using biophysical and biochemical methods. We have used computational sequence analysis tools to predict probable functions of CBP2. Based on these predictions, we have analysed its nucleic acid and ATP binding properties using recombinant cytoplasmic domain of CBP2 (rcCBP2). We have also studied its interaction with MC resident protein, PfSBP1 and studied the impact of ATP binding on the PfSBP1-CBP2 interaction. Our results highlight that CBP2 not only engages endothelial cells via its extracellular domain but is also likely to perform other important roles inside iRBCs and EVs.

## 2. Results

### 2.1 Domain analysis

CBP2 is a PEXEL positive multi-transmembrane protein that interacts with CX3CL1 receptor expressed on host endothelial cells [12]. Sequence analysis of this protein using TMHMM predicts three transmembrane regions spanning 4-26, 164-187 and 201-224 amino acid residues, while its PEXEL motif is present on residues 46-50 (RSLAE) (Fig. 1A). This motif is likely to be cleaved during export leading to formation of a 2TM protein product that finally localizes at iRBC membrane [12]. The extracellular region encompassing amino acids 188-199 is reported to bind with the chemokine CX3CL1 (Fig. 1B) [12]. Domain analysis using Pfam (v. 27) suggested that CBP2 harbours three domains, TROVE (28-161 amino acids), claudin 2 (22-244 amino acids) and LrgA (168-242 amino acids) (Fig. 1A). The TROVE domain (Telomerase, Ro and Vault) is a module of ∼300-500 residues that is found in protein components of ribonucleoprotein particles and possesses RNA binding properties [25], while LrgA is believed to transport murein hydrolases [26]. Claudin 2 is a family of proteins present in tight junctions responsible for controlling paracellular flow of molecules across epithelial layers of cells [27]. Motif scan analysis shows CBP2 to contain another small domain ‘cold shock DNA binding domain’ (37-55 residues). Additionally, the conserved domain database (CDD, NCBI) predicts a segment (161-231 amino acids) from the permease family of proteins, whose members are integral membrane proteins with six transmembrane regions. CBP2 shows 32% sequence identity with CBP1 (Fig. 1C).

**Fig. 1.**
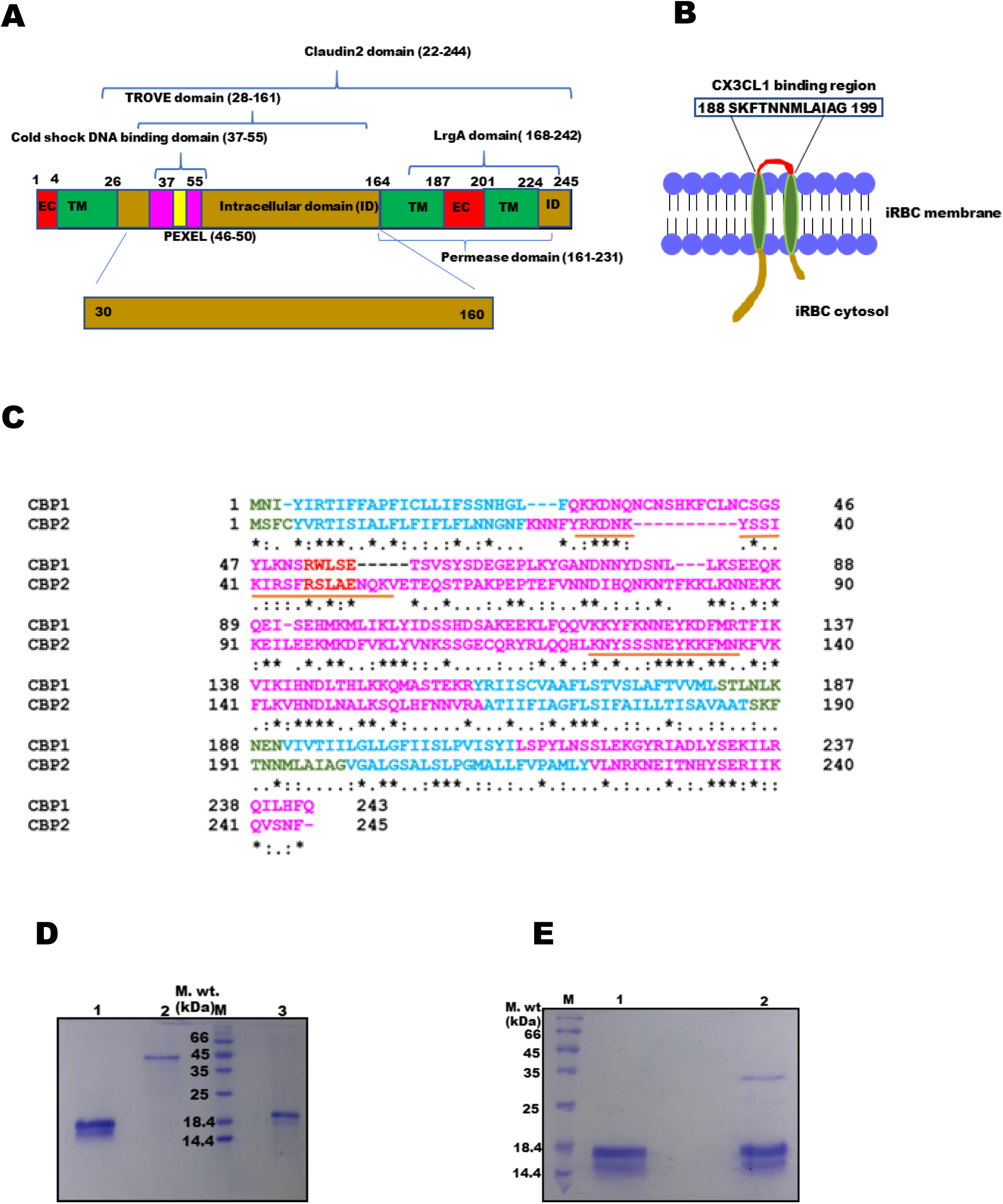
Sequence analysis of CBP2 and purification of recombinant proteins: (A) *Domain organization of CBP2*. Figure shows transmembrane regions (TM, green), extracellular regions (EC, red), PEXEL motif (PEXEL, yellow), intracellular domains (ID, brown). CBP2 harbours cold shock DNA binding domain (37-55), TROVE domain (28-161), Claudin2 domain (22-244) LrgA domain (168-242) and permease domain (161-231). Cloned region (30-160) is shown at the bottom of the panel. (B) *iRBC membrane topology of CBP2*. Figure highlights CX3CL1 binding residues (red). (C). *Pairwise sequence alignment of CBP1 and CBP2*. Residues corresponding to EC are marked in green, ID in pink, TM in blue and PEXEL in red. Nucleic acid binding residues are underlined. * represents conserved residues, : represents semiconserved residues and . represents similar residues. (D) *SDS-PAGE of purified recombinant proteins*. Lane1: rcCBP2, lane 2: ATS of PfEMP1, lane M: protein molecular weight marker, lane 3: PfSBP1. (E). *SDS-PAGE of rcCBP2 under reducing and non-reducing conditions*. Lane M: molecular weight marker, lane 1: rcCBP2 (reducing), lane 2: rcCBP2 (non-reducing).

### 2.2 Cloning, expression, and purification of rcCBP2

The DNA segment corresponding to the cytoplasmic domain of CBP2 (30 to 160 amino acid residues) was cloned in pET-28a (+) expression vector and expressed in BL21 (DE3) *E. coli* cells. The recombinant protein was purified in soluble form using HIS-Select HF Nickel Affinity and gel permeation chromatographies (Fig. 1D, lane 1), and the identity was confirmed using anti-hexa histidine antibodies by western blotting (Fig. S1A). Recombinant ATS domain of PfEMP1 (PF08_0141) and PfSBP1 were expressed and purified using various chromatographies as described earlier by Kumar *et al* (Fig. 1D, lanes 2-3) [28]. Polyclonal antibodies against purified CBP2 were commercially raised in New Zealand White rabbits. Antibody reactivity was confirmed *in vivo* by western blot analysis on iRBC extracts (iRBC pellet and supernatant after saponin lysis) from cultured Pf3D7 parasites (Fig. S1B). Antibody specificity was verified using western blot analysis on induced and uninduced *E. coli* cell lysates, where a single band was detected in the induced sample only (Fig. S1C). Also, no bands were observed in uninfected RBC lysates, confirming specificity of the raised antisera (Fig. S1B).

#### 2.3.1 rcCBP2 assumes different oligomeric states under reducing and non-reducing conditions

CBP2 has been previously shown to exist in multiple oligomeric states (monomer, dimer and trimer) in iRBC membrane fraction when probed with antibodies against CBP2 peptide [12]. Since CBP2 contains cysteine residues (C4 & C113), we separated rcCBP2 on SDS-PAGE under reducing and non-reducing conditions. Here, a band corresponding to the size of its monomeric state (∼ 21 kDa) was observed in the reducing gel, while an additional prominent band corresponding to its dimeric state was also visible under non-reducing conditions (Fig. 1E).

#### 2.3.2 rcCBP2 binds with ATP

Our domain analysis and experimental data reveal that CBP2 is a 2TM protein with inter-chain disulphide linkages. Therefore, we tested ATP binding of rcCBP2 to understand if CBP2 behaves as a P2X purinergic receptor, which are 2TM ligand gated ion channel proteins that bind with ATP and contain disulfide bonds [29]. Interaction of rcCBP2 with ATP was studied using native-PAGE, dynamic light scattering (DLS) and nuclear magnetic resonance (NMR) spectroscopy.

Native-PAGE analysis was performed on rcCBP2 (with and without ATP) followed by immunoblotting with anti-hexa histidine antibodies. Multiple bands with reduced mobility were observed upon incubation with ATP, suggestive of changes in the structure or oligomeric state of rcCBP2 (Fig. 2A). Dynamic light scattering (DLS) further verified the interaction of rcCBP2 with ATP leading to structural changes as indicated by the increased hydrodynamic diameter of the protein in solution. The average hydrodynamic diameters increased from 133.5 nm and 334.3 nm (without ATP) to 533.4 nm in the presence of ATP (Fig. 2B).

**Fig. 2.**
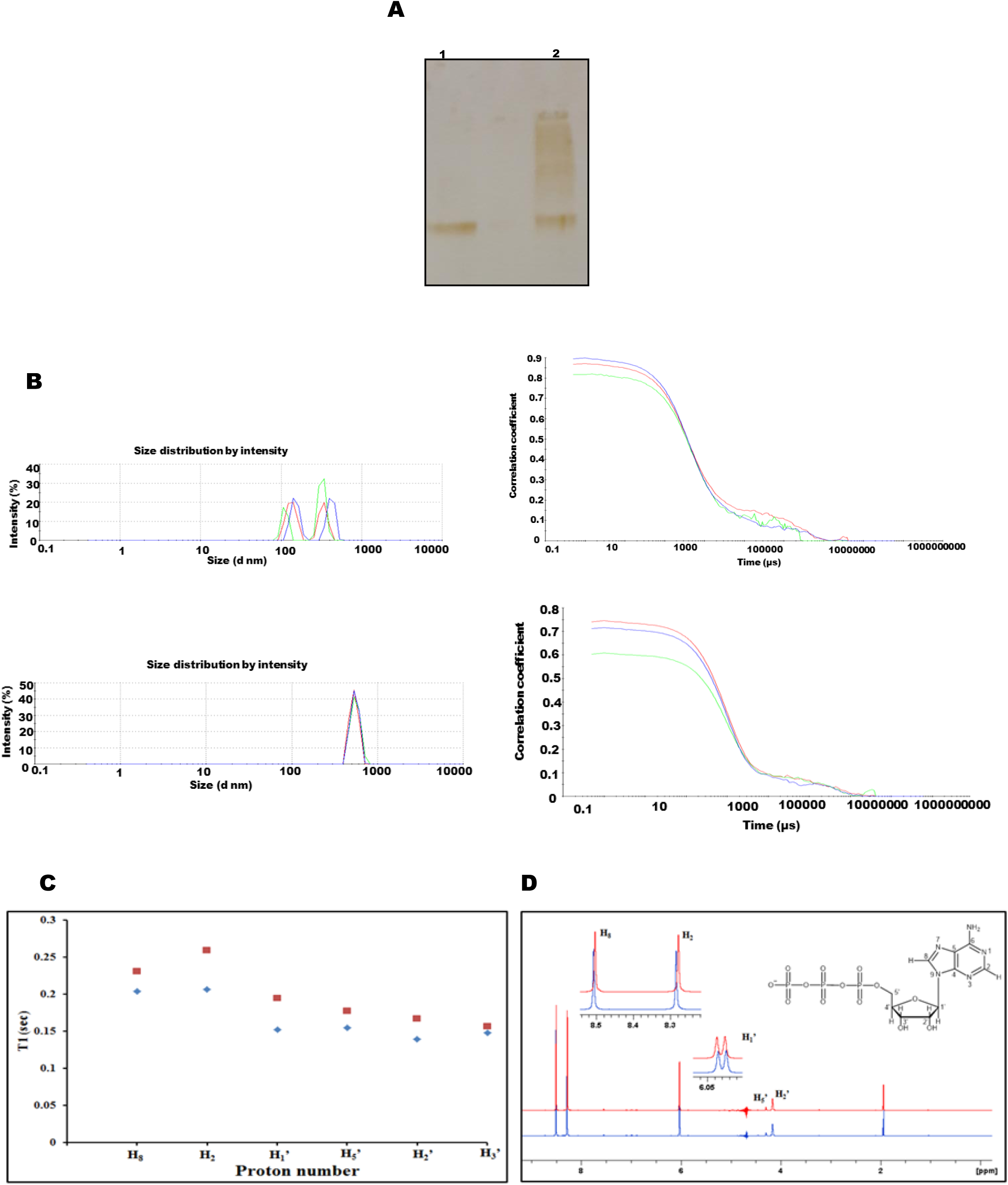
rcCBP2-ATP interaction: **(A)** *Immunoblot of native-PAGE showing rcCBP2-ATP interaction*. rcCBP2 was run on 15% native-PAGE and subjected to immunoblotting using anti-hexa histidine-HRP conjugated antibodies (1:2000). Lane 1: rcCBP2, lane 2: rcCBP2 + ATP (B) *DLS showing rcCBP2-ATP binding*. DLS of rcCBP2 (top panel) and rcCBP2 + ATP (bottom panel). Size distribution curves of rcCBP2 (by intensity) (left panel); correlograms corresponding to size distribution curves (right panel). (C, D) *NMR analysis of ATP to study structural changes upon rcCBP2 binding*. (C) ^1^H T_1_ relaxation times of ATP (40 mM) in the absence (red dots) and presence (blue dots) of rcCBP2. (D) ^1^H NMR spectrum of ATP (blue trace) and ATP in the presence of protein (red trace).

#### 2.3.3 H2 and H8 of adenine directly interact with rcCBP2

Interaction of ATP with rcCBP2 at atomic level was further analysed by spin-lattice relaxation times and ^1^H NMR chemical shift measurements. Spin-lattice relaxation time (T_1_) values were measured on the solution of ATP in the absence and presence of protein. Addition of rcCBP2 (5 µl of 1 mg/ml) to a solution of ATP resulted in decrease in spin-lattice relaxation time (T_1_) as the motion of ATP slowed down on binding with the protein (Fig. 2C). It is observed that T_1_ of H2 and H8 of adenine and anomeric proton (H’-1) of sugar unit exhibited considerable decrease upon addition of protein. Thus, the T_1_ measurement results indicate that ATP has significant intermolecular interactions with the protein. ^1^H NMR spectrum of ATP (40 mM) was recorded in 0.5 ml D_2_O: H_2_O (90:10) at 25 °C (blue trace, Fig. 2D). Addition of 5 µl of protein to the solution of ATP resulted in significant upfield shift of H-2, H-8, H’-1 whereas no visible change in the CH protons of sugar unit was observed. Hence, these results confirm the atomic level interaction of ATP with rcCBP2.

### 2.4 rcCBP2 binds with nucleic acids

The cytoplasmic domain of CBP2 is predicted to carry nucleic acid binding domains (Fig. 1A). Therefore, we used gel retardation assays and pull-down assays to test the nucleic acid binding properties of rcCBP2. rcCBP2 was incubated separately with ss-DNA oligomers, ds-DNA (plasmid) and RNA, and subjected to agarose gel electrophoresis. Retardation in migration of nucleic acids was observed suggesting that CBP2 binds both with DNA and RNA (Fig. 3A & B). A shift in rcCBP2 mobility could also be observed when these samples were run on native-PAGE, transferred to NC membrane and probed using protein specific antibodies (Fig. 3C). Negative controls (PfJ23, PfSBP1 and ATS of PfEMP1) did not show any shift in an agarose gel upon incubation with ss-DNA oligomers, highlighting the specificity of rcCBP2-nucleic acid interaction (Fig. 3D).

**Fig. 3.**
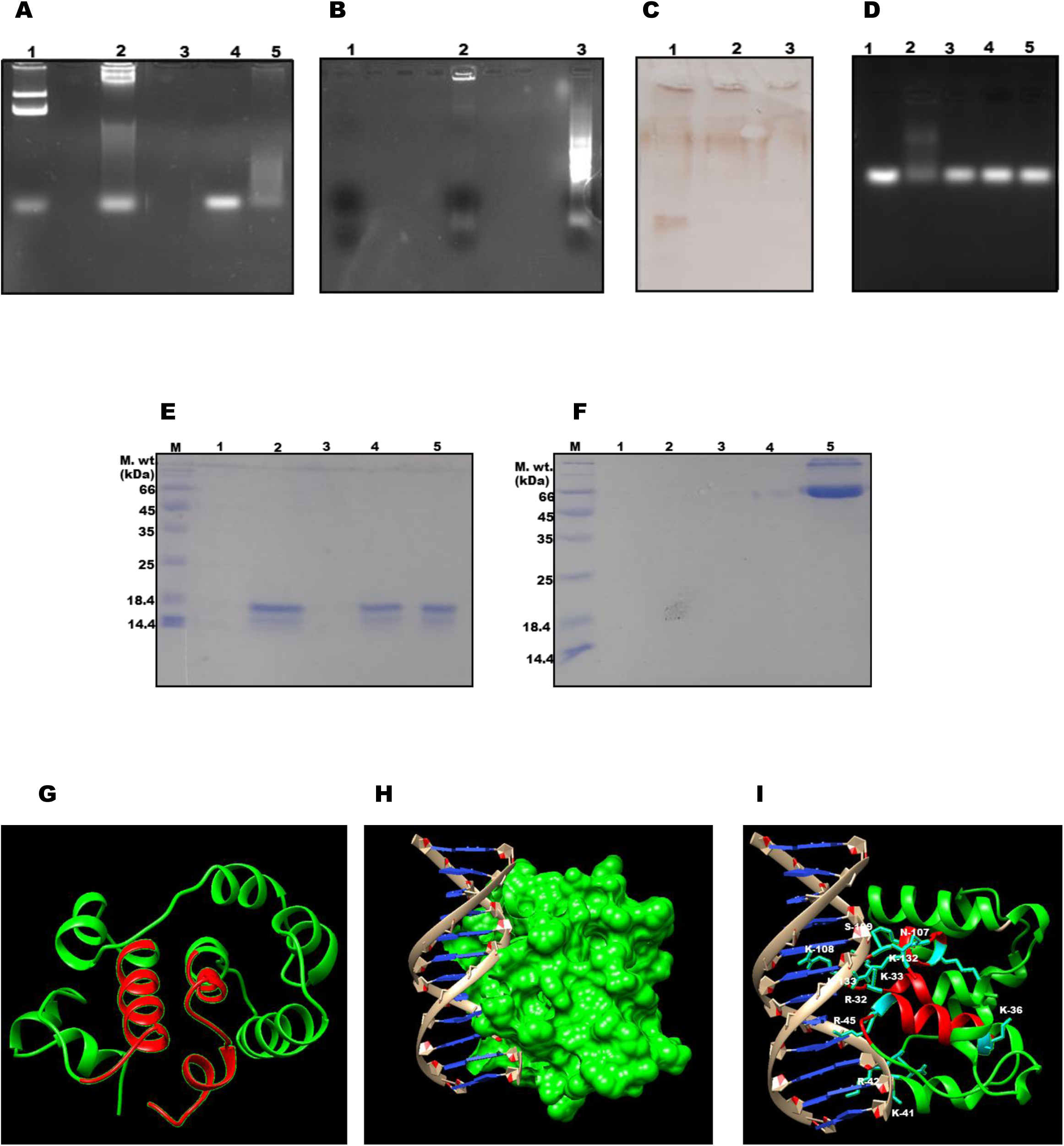
rcCBP2-nucleic acid interaction: (A-D) *Gel retardation assays showing specific rcCBP2-nucleic acid interaction*. (A) Agarose gel showing rcCBP2-DNA binding. Lane 1: ds-DNA (plasmid), lane 2: rcCBP2+ds-DNA, lane 3: rcCBP2, lane 4: ss-DNA oligomers, lane 5: ss-DNA oligomers + rcCBP2 (B) Agarose gel showing rcCBP2-RNA binding. Lane 1: rcCBP2, lane 2: rcCBP2 + RNA, lane 3: RNA. (C) Immunoblot showing specificity of rcCBP2-mucleic acid interaction. 15% native-PAGE probed using anti-CBP2 antibodies; Lane 1: rcCBP2, lane 2: rcCBP2+ds-DNA, lane 3: rcCBP2+ ss-DNA oligomers. (D) Agarose gel showing specificity of rcCBP2 binding with ss-DNA oligomers. Lane1: ss-DNA oligomers, lane 2: rcCBP2 + ss-DNA oligomers, lane 3: PfJ23 + ss DNA oligomers, lane 4: PfSBP1 + ss DNA oligomers, lane 5: ATS of PfEMP1 + ss DNA oligomers. (E-F) *rcCBP2-DNA interaction using pull down assay*. rcCBP2(E) and negative control (BSA, F) were incubated with ss-DNA oligomer/ ds-DNA immobilized cellulose beads in a pull-down assay and loaded on 15% SDS-PAGE. Lane M: protein molecular weight marker, lane 1: ss-DNA oligomer wash, lane 2: ss-DNA oligomer + rcCBP2, lane 3:ds-DNA wash, lane 4: ds-DNA + rcCBP2, lane 5: rcCBP2/ BSA(G) *Three-dimensional structure of rcCBP2 generated using homology modeling*. Ribbon diagram of rcCBP2 (green) with predicted nucleic acid binding residues highlighted in red. (H) *rcCBP2-ds-DNA docked complex*. rcCBP2 is shown in a space fill form (green) and DNA in brown (sugar-phosphate backbone) and blue (H-bonded nucleotide bases). (I) *rcCBP2-ds-DNA docked complex highlighting residues of rcCBP2 at the interaction interface*. rcCBP2 is shown in ribbon format, DNA in brown (sugar-phosphate backbone) and blue (H-bonded nucleotide bases), and the interacting residues are represented as ball and sticks (cyan).

The nucleic acid binding property of rcCBP2 was also analysed using nucleic acid hybridization assay with cellulose beads containing immobilized calf thymus ss-DNA and ds-DNA (Fig. 3E). rcCBP2 showed binding with both ss-DNA and ds-DNA, while no binding was observed in the negative control (BSA) (Fig. 3F). These results together highlight that the cytoplasmic domain of CBP2 has nucleic acid binding properties.

### 2.5 Structure prediction of rcCBP2 and molecular docking with DNA

Homology modelling of rcCBP2 was performed using the crystal structure of native 8-oxoguanine DNA glycosylase from *Pyrobaculum aerophilum* (PDB ID: 1XQO) as a template. The generated models (amino acid residues 30 to 160 of CBP2) were validated by using ERRAT and RAMPAGE, and the best model was used for further analysis (Fig. 3G). Scores obtained for the best model are present in Fig. S2.

Docking of the modelled CBP2 with B-DNA (PDB ID: 1BNA) was performed using Patchdock at residues 30-50 and 122-136 which were predicted to bind DNA by Motif Scan. These two stretches of amino acid residues are helical and found to be embedded in major groove of B-DNA (Fig. 3H). Apart from basic amino acids (R-32, K-33, K-36, K-41, R-42 R-45, K-108, K-123, K-132, K-133 and K-136), residues N-107, S-109, F-137 and V-138 are also involved in CBP2-DNA interaction (Fig. 3I).

#### 2.6.1 rcCBP2 interacts directly with PfSBP1

CBP2 has been reported to co-localize with MC resident protein PfSBP1 at the ring stage of parasite development [14]. Therefore, we carried out PfSBP1-rcCBP2 interaction studies using dot-blot assay. PfSBP1, negative controls (ATS of PfEMP1 and BSA) and positive control (rcCBP2) were immobilized on NC membrane, incubated with rcCBP2 and probed with anti-CBP2 antibodies. Signals were observed in the positive control and the spot containing PfSBP1, suggesting interaction between rcCBP2 and PfSBP1. No signal could be seen in the negative controls (Fig. 4A), or an identical blot probed with pre-immune sera (Fig. S2). Cross-reactivity of PfSBP1 with anti-CBP2 antibodies was ruled out using another blot where PfSBP1 was spotted and probed with anti-CBP2 antibodies (Fig. S2). Specificity of the rcCBP2-PfSBP1 interaction was checked by using native-PAGE followed by immunoblotting. Our results showed a clear shift in the band corresponding to rcCBP2 upon incubation of the protein with PfSBP1 (Fig. 4B).

**Fig. 4.**
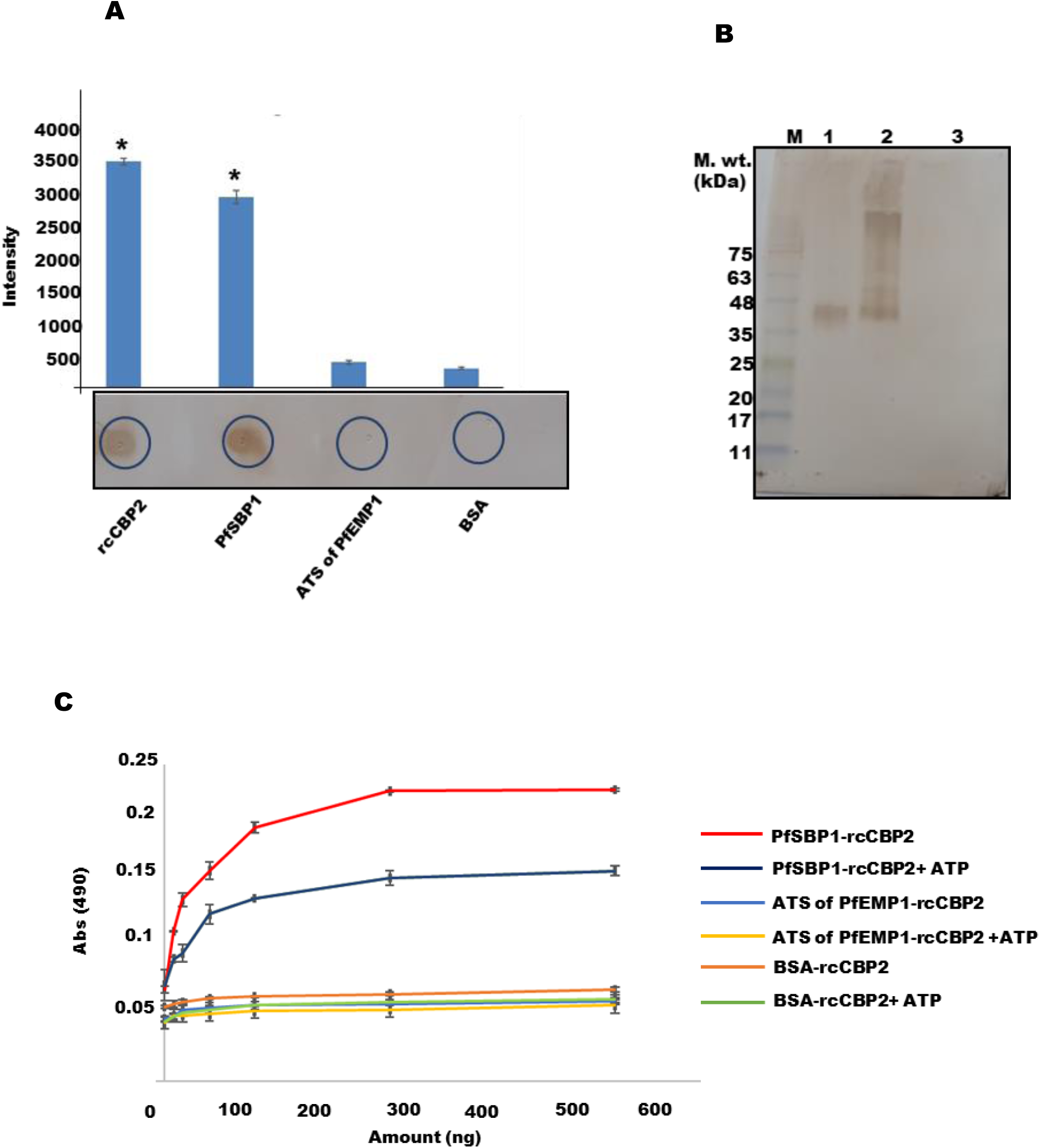
rcCBP2-PfSBP1 interaction: (A) *Dot-blot assay*. Proteins were spotted as labelled on the blot, incubated with rcCBP2 and probed with anti CBP2 antibodies (Bottom panel). Dot intensities were measured and plotted as a bar chart (Top panel). (B) *Immunoblot showing specificity of rcCBP2-PfSBP1 interaction*. 15% native-PAGE probed with anti-CBP2 antibodies. Lane 1: ladder, lane 2: rcCBP2, lane 3: rcCBP2+ PfSBP1, lane 4: PfSBP1. (C) *Effect of ATP on rcCBP2-PfSBP1 interaction using indirect ELISA*. PfSBP1 (experimental) and negative controls (ATS of PfEMP1 or BSA) were on the wells, incubated with rcCBP2 (with or without ATP) and probed with anti CBP2 antibodies. Experiments were performed in triplicates and concentration dependent binding curves were plotted as labeled on the figure.

#### 2.6.2 ATP binding to CBP2 weakens the interaction of CBP2 and PfSBP1

Interaction between rcCBP2 and PfSBP1 was further verified by semi quantitative indirect ELISA, where PfSBP1 was coated in the wells, followed by incubation with increasing amounts of rcCBP2 (10 ng to 500 ng). Increase in absorbance was observed with increasing amount of rcCBP2 confirming the interaction between these proteins (Fig. 4C). Interestingly, ATP binding of rcCBP2 weakened its interaction with PfSBP1 (Fig. 4C) possibly due to structural alterations in rcCBP2 structure upon ATP binding. Negative controls (BSA and ATS of PfEMP1) did not show any binding to rcCBP2 both in its ATP bound and unbound forms, highlighting the specificity of the interaction.

## 3. Discussion

Sequence analysis of CBP2 predicts this PEXEL positive protein to contain three transmembrane (TM) regions along with a pair of cytoplasmic domains and short extracellular regions each (Fig.1A). The extracellular loop sandwiched between the second and third TM regions is reported to bind with membrane associated CX3CL1 on endothelial cells of microvessels [12]. Expression of CX3CL1 was also observed at the site of iRBC sequestration in a cerebral malaria patient [12], suggestive of the involvement of CBPs in cytoadherence. In the present study, we have cloned, expressed and characterized the unexplored N-terminal cytoplasmic region (30-160) of CBP2 which contains a cold shock DNA binding domain (Fig. 1A). Migration of rcCBP2 under reducing and non-reducing condition revealed the involvement of disulphide bonds in its oligomerization (Fig. 1D).

Our domain analysis (Fig. 1A) predicts CBP2 to contain two TM regions after cleavage of the PEXEL motif by Plasmepsin V [14]. Based on the presence of two TM domains and disulphide bonds in processed CBP2 expressed on the iRBC membrane, this protein is likely to act as a P2X receptor. P2X receptors are purinergic ligand gated ion channels that get activated by ATP and other nucleotides [30-31]. These receptors are 2TM proteins that may have disulphide linkages and show ATP binding to mediate various biological processes [32]. Erythrocytes respond to various stimuli by releasing ATP extracellularly, which is known to activate purinergic receptors and play a role in erythrocyte invasion by parasites [33-34]. Therefore, we investigated the binding of rcCBP2 with ATP using native-PAGE, DLS and NMR spectroscopy (Fig. 2A, B, C & D). Our results show that ATP not only binds rcCBP2, but also induces changes in the structure of this protein. Therefore, we hypothesize that CBP2 may have a role in transmission of signals in iRBCs similar to P2X receptors.

Proteomics analysis has shown presence of CBP2, but not CBP1 in iRBC derived microvesicles (MVs) [20-21]. MVs are small vesicles that are released from various cell types to affect physiological processes [35]. Infection with *Plasmodium* species is known to cause elevated release of microvesicles from iRBCs, which fuse with host cells to cause immunomodulation and gametocytogenesis [20]. Interestingly, delivery of these MVs to endothelial cells (ECs) is responsible for altered gene expression and reduced barrier function in recipient ECs through host miRNAs packaged in MVs [36]. iRBC generated MVs are also loaded with various other important molecules like parasite antigens, parasite mRNAs, Y-RNA, vault RNA and other non-coding RNAs for delivery to ECs [37-38]. Our current study depicts binding of rcCBP2 (cytoplasmic domain) with different nucleic acids (DNA and RNA) using gel retardation (Fig. 3A, B & C) and pull-down assays (Fig. 3D & E). Our *in silico* domain analysis of CBP2 predicts the presence of a TROVE module (telomerase, Ro and vault), which is a common element of ribonucleoprotein particles (RNPs) [25]. Ro protein interacts with Y RNAs and is also known to be involved in autoimmunity [39]. Vault is a high molecular weight protein that binds vault RNA and has a suggested role in multidrug resistance [40]. Therefore, the cytoplasmic domain of CBP2 is very likely to bind different RNAs within MVs and /or in iRBC cytosol. Based on interaction of extracellular domain of CBP2 with CX3CL1 and its cytoplasmic domain with nucleic acids, we hypothesize that CBP2 may serve to target MVs to ECs while delivering cargo (different types of RNAs) inside MVs bound to its cytosolic domain (Fig. 5).

**Fig. 5.**
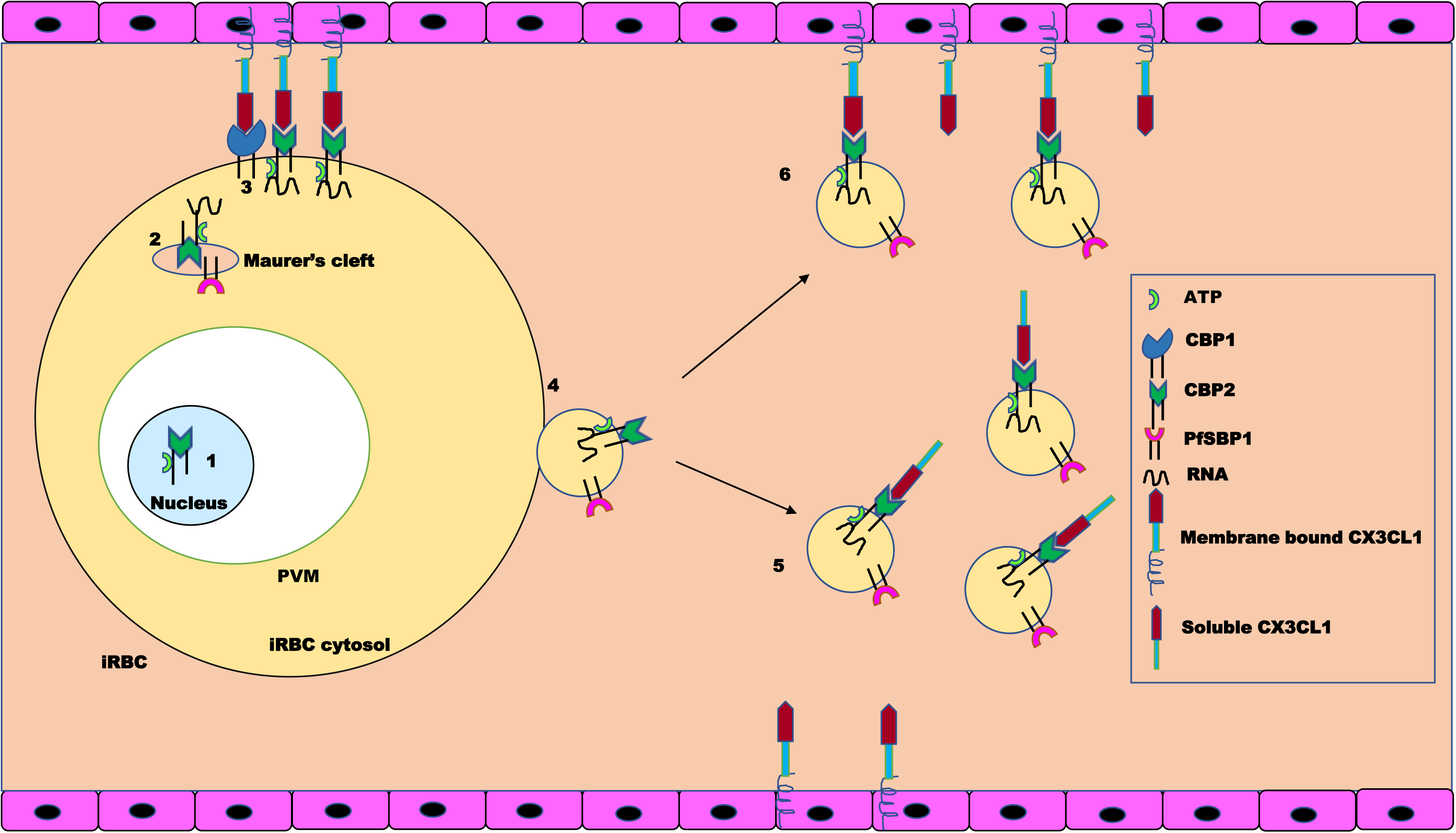
Hypothetical model for probable function of CBP2: Step 1: CBP2 localizes to the parasite nucleus in early ring stage [13]. Step 2: After its translocation to iRBCs, it co-localizes with PfSBP1 in MCs [14] Step 3: CBP1 and CBP2 are exported to the iRBC membrane where they mediate cytoadherence of iRBCs by interaction with CX3CL1 on endothelial cells [12] Step 4: EVs containing CBP2 along with other host and parasite RNAs and proteins (e.g. PfSBP1) are released from iRBCs. Cytoplasmic domain of CBP2 may carry bound host and parasite RNAs from iRBC cytosol to EVs for delivery to ECs. CBP2 incorporated in EVs binds to the chemokine domain of soluble CX3CL1 (Step 5) and membrane bound CX3CL1 (Step 6) to deliver cargo to host endothelial cells. ATP binding of CBP2 is likely at any of these steps.

A report has shown co-localization of CBP2 with PfSBP1 in a punctuate pattern in ring stage iRBCs, suggestive of their presence in MCs possibly *en route* to the iRBC surface [14]. PfSBP1 is a MC resident transmembrane protein with reported roles in PfEMP1 export and anchorage with RBC membrane proteins [41]. Recent proteomics data on iRBC generated extracellular vesicles reveals the presence of several parasite proteins including MC components (PfSBP1, REX1/2, MAHRP1/2) and invasion related proteins (EBA-175, EBA-181, RhopH2/H3) etc. in EVs [20]. Co-existence of CBP2 and PfSBP1 in these compartments led us to test direct interaction between these proteins. Our dot-blot, native-PAGE and ELISA assays clearly illustrate binding between rcCBP2 and the recombinant MC luminal domain of PfSBP1 (Fig. 4A, B & C). Since both these recombinant protein stretches would assume opposite transmembrane orientation at MCs and iRBC surface, we hypothesize that interaction between the N-terminal luminal domain of PfSBP1 and cytosolic domain of CBP2 is most likely either within parasites, or during their export to MCs via the iRBC cytosol [15]. Interestingly, binding strength of the PfSBP1-rcCBP2 pair showed significant reduction in the presence of ATP, possibly due to conformational changes induced in rcCBP2 upon ATP binding. Since ATP plays an important role in transmission of signals in malaria parasites [42], alteration of protein-protein interaction dynamics in response to ATP is likely to have roles in disease pathogenesis. However, the significance and exact location of this interaction *in vivo* needs to be elucidated in parasite biology.

Based on our results and previous reports, we propose a model describing the export and probable function of CBP2 in *Plasmodium falciparum* biology (Fig. 5). CBP2 localizes to the parasite nucleus during early stages of development (Step1) [13] and is later exported to the surface of iRBCs [12] via MCs [14] (Steps 2, 3). At the surface of parasitized erythrocytes, CBP1 and 2 engage endothelial chemokine receptor CX3CL1 via their extracellular loop regions [12], while CBP2 may interact with cytosolic parasite and/ or host RNAs through its C-terminal cytoplasmic domain (Step 3). CBP2 is then packaged into extracellular vesicles carrying several other parasite antigens and parasite/ host RNAs as cargo (Step 4). These vesicles bud from iRBCs and bind CX3CL1 present on endothelial cells (Steps 5, 6). We hypothesize that delivery of EVs and its cargo to ECs may involve CBP2. In a nutshell, we suggest a key role for CBP2 in packaging and delivery of parasite and erythrocytic RNAs to host endothelial cells for their modification. Our findings have important implications in understanding malaria pathogenesis and host-pathogen interactions.

## 4. Materials and methods

### 4.1 Bioinformatics analysis

Sequence of CBP2 (PF3D7_1301700) was retrieved from PlasmoDB [43], and domain analysis was performed using Pfam version 27 [44] and NCBI conserved domain database (CDD) [45]. Transmembrane regions were predicted by using TMHMM [46], while pairwise sequence alignment of CBP1 & 2 proteins was performed using EMBOSS Needle tool (EMBL-EBI) [47].

### 4.2 Parasite culture

*Plasmodium falciparum* 3D7 was cultured in RPMI 1640 supplemented (HiMedia) with heat inactivated 10% human serum using O^+^ RBC’s in the presence of 90 % N_2_, 5 % O_2_ and 5 % CO_2_. Cultured parasites were maintained at 5% hematocrit and 5% parasitemia.

### 4.3 Cloning, expression, purification of rcCBP2 and generation of specific antisera

The DNA fragment corresponding to the cytoplasmic domain of CBP2 (30 to 160 amino acid residues) was PCR amplified using Pf3D7 genomic DNA as a template, cloned into pET28a plasmid (Novagen, Merck KGa, Madison, WI, USA), and expressed in *E. coli* BL21(DE3) cells. Purification of rcCBP2 was performed by HIS-Select HF Nickel Affinity (Sigma) and gel permeation chromatography on Superdex 75 column (GE Healthcare Life Sciences). Purified rcCBP2 was used for raising specific antisera in rabbits commercially (Radiant research, Bangalore, India). The effect of disulphide linkages on the oligomeric state of rcCBP2 was checked by running the protein on SDS-PAGE under reducing and non-reducing conditions.

### 4.4 Immunoblotting using purified protein and parasite lysates

Recombinant rcCBP2 was run on SDS-PAGE and subjected to western blotting using anti-hexa histidine HRP (horse radish peroxidase) conjugated antibodies (Sigma; 1:2000) to confirm identity of the purified protein.

*In vivo* protein expression was checked in mixed stage saponin lysed parasite cultures of Pf3D7 (parasitemia 5-10%). Here, the pellet was extensively washed with 1X PBS to remove any residual cytosolic contents. Parasite pellet, supernatant from lysate and purified recombinant rcCBP2 (positive control) along with pellet and supernatant of uninfected RBCs were resolved on 15% SDS-PAGE and transferred to nitrocellulose (NC) membrane to perform western blot analysis using protein specific antibodies (1:4000) as described above.

Antibody validation was performed by testing the raised polyclonal antibodies on uninduced and induced *E. coli* cell lysates (transformed with recombinant plasmid) in a western blot experiment. 5% BSA in 1X PBS was used for blocking overnight at 4^□^C and probed with rabbit anti-CBP2 (1:5000) antisera followed by HRP conjugated goat anti rabbit IgG antibodies (1:2000). All blots were developed using 3,3’-diaminobenzidine (DAB)/H_2_O_2_ substrate.

### 4.5 Effect of ATP on rcCBP2

#### 4.5.1 Native-PAGE

Interaction of rcCBP2 with ATP was assessed using Native-PAGE. Recombinant protein (5 µg) was incubated with 10 µM ATP in binding buffer (20 mM tris Cl pH 8, 2.5mM MgCl_2_, 2.5mM MnCl_2_, 1mM sodium orthovanadate, 0.5mM sodium fluoride) at 30°C for 1.5 hrs and run on 15% native-PAGE. The gel was transferred on nitrocellulose (NC) membrane, probed using monoclonal anti-hexa histidine HRP conjugated antibodies and developed.

#### 4.5.2 Dynamic light scattering

Purified rcCBP2 was studied by dynamic light scattering (DLS) (Zetasizer Nano ZS, Malvern instrument). Size measurements were carried out after incubating solutions of rcCBP2 (1µM) in the absence and presence of 5µM ATP in binding buffer for 1.5 hours. Three scans were performed for each sample at 25°C.

#### 4.5.3 ^1^H NMR T_1_ relaxation time measurements

Interaction of ATP and CBP2 was also analysed by ^1^H nuclear magnetic resonance (NMR). ^1^H NMR T_1_ relaxation time measurements were performed on Bruker Avance 500 NMR spectrometer at 298.2 K. The longitudinal relaxation time (T_1_) was determined by 180°-90° inversion recovery pulse sequence. 16 values of delay time (τ) were applied and 16 scans for each τ value were recorded. The pre-acquisition delay (D1) was set to 2 × T_1_ (5 s) of the longest relaxation time. The value of the longitudinal relaxation time was obtained with the help of T1/T2 relaxation module of Topspin as described in the manual of this software, whereas the fitting function “invec” and fitting type ‘‘area” was used.

### 4.6 *In vitro* nucleic acid binding

#### 4.6.1 Gel retardation assays

1µg of purified rcCBP2 was incubated with 100 ng of DNA [single stranded (ss) DNA oligomers/ plasmid] or RNA (total Pf RNA isolated as described in Moll *et al* [48]) in 2X nucleic acid binding buffer (3 mM MgCl_2_, 50% glycerol, 1M HEPES, 100 mM DTT, 1M tris-Cl pH 8) at 37°C for 1 hr. Migration patterns of nucleic acid samples incubated with rcCBP2 were studied by agarose gels stained with ethidium bromide. Specificity of the interaction of rcCBP2 with nucleic acids was ascertained by incubating ss DNA oligomers with other proteins i.e. BR5 domain of PfSBP1, ATS of PfEMP1 (PF08_0141) and PfJ23 before running the mixtures on an agarose gel. These proteins were purified according to protocols described in Kumar *et al* [28] and Kaur *et al* [49].

rcCBP2 incubated with DNA (plasmid/ ss-DNA oligomers) was also run on 15% native-PAGE and transferred on NC membrane. Identity of rcCBP2 was confirmed by western blot analysis using protein specific antibodies (1:5000). The blot was probed and developed as explained above.

#### 4.6.2 Bead based rcCBP2-DNA pull down assay

The nucleic acid binding property of rcCBP2 was further validated using cellulose beads with immobilized single and double stranded calf thymus DNA (Sigma Aldrich, USA). 3µg of rcCBP2 was mixed with 500 µl cellulose matrix [1 mg/ml matrix prepared in 50 mM sodium phosphate (pH 7.6), 150 mM NaCl] and incubated at 4°C for 15 min. After centrifugation at 3000 rpm for 1 min at 4°C, the supernatant was discarded, and the bead pellet was washed five times with 200 µl buffer. The washed beads were boiled in 1X SDS-PAGE sample loading dye before loading on 15% SDS-PAGE. BSA was used as negative control in this assay [50].

#### 4.6.3 Structure prediction, validation and molecular docking

Three-dimensional structure of CBP2 was predicted using homology modeling where PDB ID: 1XQO (crystal structure of 8-oxoguanine DNA glycosylase from *Pyrobaculum aerophilum*) was used as the template [51]. The basic modeling approach of Modeller 9.14 [52] was used for structure prediction, and the predicted models were validated using ERRAT [53] and RAMPAGE [54]. UCSF Chimera was used for visualization of structures [55].

Docking analysis between CBP2 and B-DNA (PDB ID: 1BNA) was carried out using the Patchdock server v. beta 1.3 [56-57]. The default parameters of Patchdock web-server were used for docking, and RMSD (root mean square deviation) value of 1.5 was used for clustering.

### 4.7 *In vitro* interaction studies between rcCBP2 and PfSBP1

#### 4.7.1 Dot-blot assay

*In vitro* binding of rcCBP2 and PfSBP1 was studied using dot blot assays. Different proteins (1µg each) i.e. rcCBP2 (positive control), BR5 domain of PfSBP1 [48], and negative controls (ATS domain of PfEMP1 [49] and BSA) were immobilized on NC membrane and blocked with 5% BSA in 1X PBS for 2 hrs at room temperature (RT). After blocking, blots were washed with 1X PBS and probed with rcCBP2 (20 µg/ 5ml) for 1hr at RT. These blots were washed twice each with PBST (PBS with 0.025% tween 20) and PBS, and then incubated for 1hr with anti-CBP2 antibodies (1:5000) followed by anti-rabbit HRP conjugated secondary antibodies (1:2000) before developing with DAB/H_2_O_2_ chromogenic substrate. BR-5 region of PfSBP1 and ATS domain of PfEMP1 were expressed and purified as described by Kumar *et al* [28]. Densitometric analysis of the blots was performed using ImageJ [58], and statistical analysis (One-way ANOVA) was done using SigmaSTAT [59].

#### 4.7.2. Native-PAGE and immunoblotting

5 µg each of purified rcCBP2 and PfSBP1 were incubated together at 37°C for 30 min. This reaction mixture containing rcCBP2 and PfSBP1 was run on 15% native-PAGE, transferred to NC membrane and blocked with 5% BSA. The blot was probed with anti-CBP2 antibodies and developed as described above.

#### 4.7.3 Plate based protein-protein interaction studies

100ng each of purified PfSBP1 and negative controls (BSA and ATS of PfEMP1) were coated on a 96 well ELISA plate before blocking with 5% BSA in 1X PBS at 4°C overnight. The coated ligands were incubated with increasing amounts of rcCBP2 (10, 20, 50, 100, 250 and 500 ng) for 2 hr at RT followed by washing with PBST and PBS (2 washes each). The plate was incubated with anti-CBP2 (1:5000) antibodies followed by goat anti-rabbit HRP conjugated secondary antibodies (1:10000) for 2 hrs each. Color was developed using 1mg/ml *o*-phenylenediamine dihydrochloride (OPD) and H_2_O_2_, the reaction stopped with 3N HCl and the absorbance recorded at 490 nm using a microplate reader (Multiskan Ascent, Thermo fisher scientific).

A similar plate-based binding assay was also performed to determine the effect of ATP binding to rcCBP2 on PfSBP1-rcCBP2 interaction. Here also, 100 ng proteins were coated on a 96 well plate and incubated with the same increasing amounts of rcCBP2 as taken above. For this assay, rcCBP2 was pre-incubated with 10 µM ATP in binding buffer at 30°C for 1.5 hours before checking its interaction with PfSBP1 and comparing it with an untreated sample. BSA and ATS of PfEMP1 were used as negative controls.

## Acknowledgements

RS is a DBT - Senior Research Fellow (DBT-SRF), while JK was a Department of Science and Technology (DST)-Inspire fellow. Laboratories of PCM and RH were funded by Department of Biotechnology (DBT), Govt. of India. We acknowledge the instrumentation facility present at Emerging Life Sciences (UGC-UPE).

## Authors’ contribution

PCM conceived the idea. RS conducted experiments. PCM, RS and RH analysed the data and wrote the manuscript. JK was involved in PfSBP1 interaction studies. VS was involved in parasite culturing. PS conducted NMR studies and analysed these data.

---

**Fig. S1.**
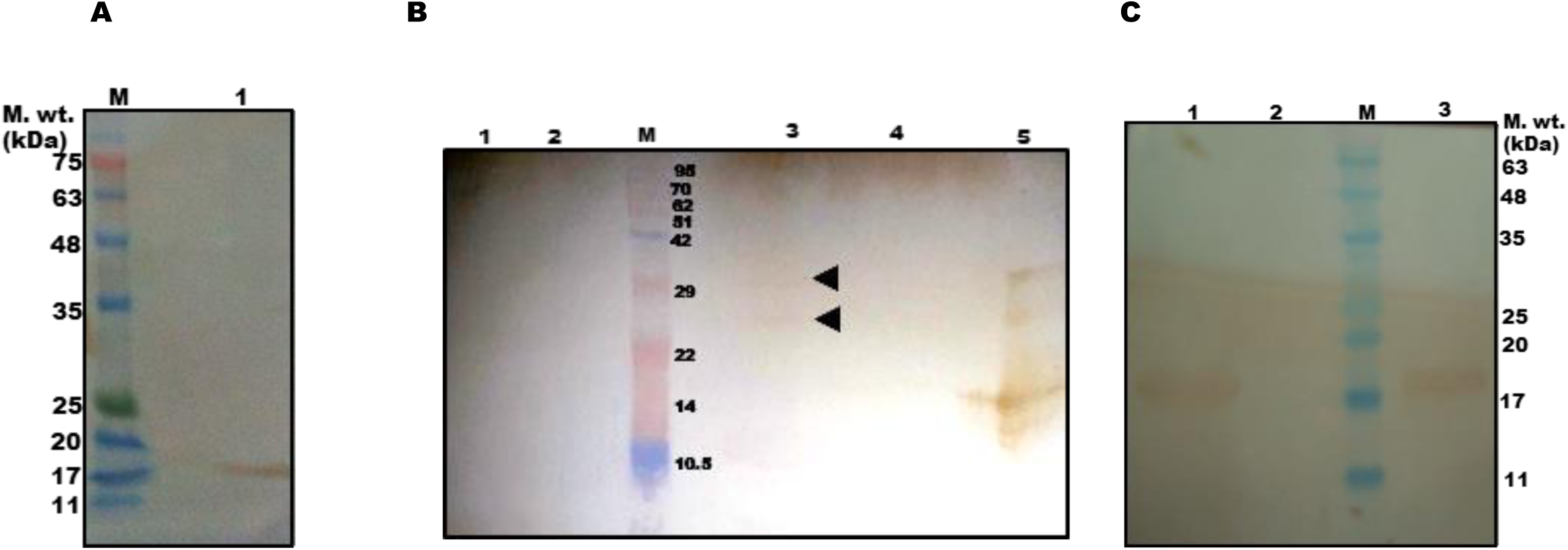
Western blots confirming identity of rcCBP2 and specificity of raised antisera: (A) *Western blot analysis of purified rcCBP2 using anti-hexa histidine antibodies*. Lane M: protein molecular weight marker, lane 1: rcCBP2. (B) *Western blot analysis of Pf3D7 parasite lysates to check in vivo expression of CBP2 using anti-CBP2 antibodies and antibody specificity of raised antisera*. Lane 1: uninfected RBC pellet, lane 2: uninfected RBC cytosol, lane M: protein molecular weight marker, lane 3: iRBC pellet, lane 4: iRBC cytosol, lane 5: rcCBP2 (positive control). (C) *Western blot analysis of E. coli lysates to assess the specificity of anti-CBP2 antibodies*. Lane 1: induced *E. coli* (BL21(DE3)) cells, lane 2: uninduced *E. coli* (BL21(DE3)) cells, lane M: protein molecular weight marker, lane 3: rcCBP2 (positive control).

**Fig. S2.**
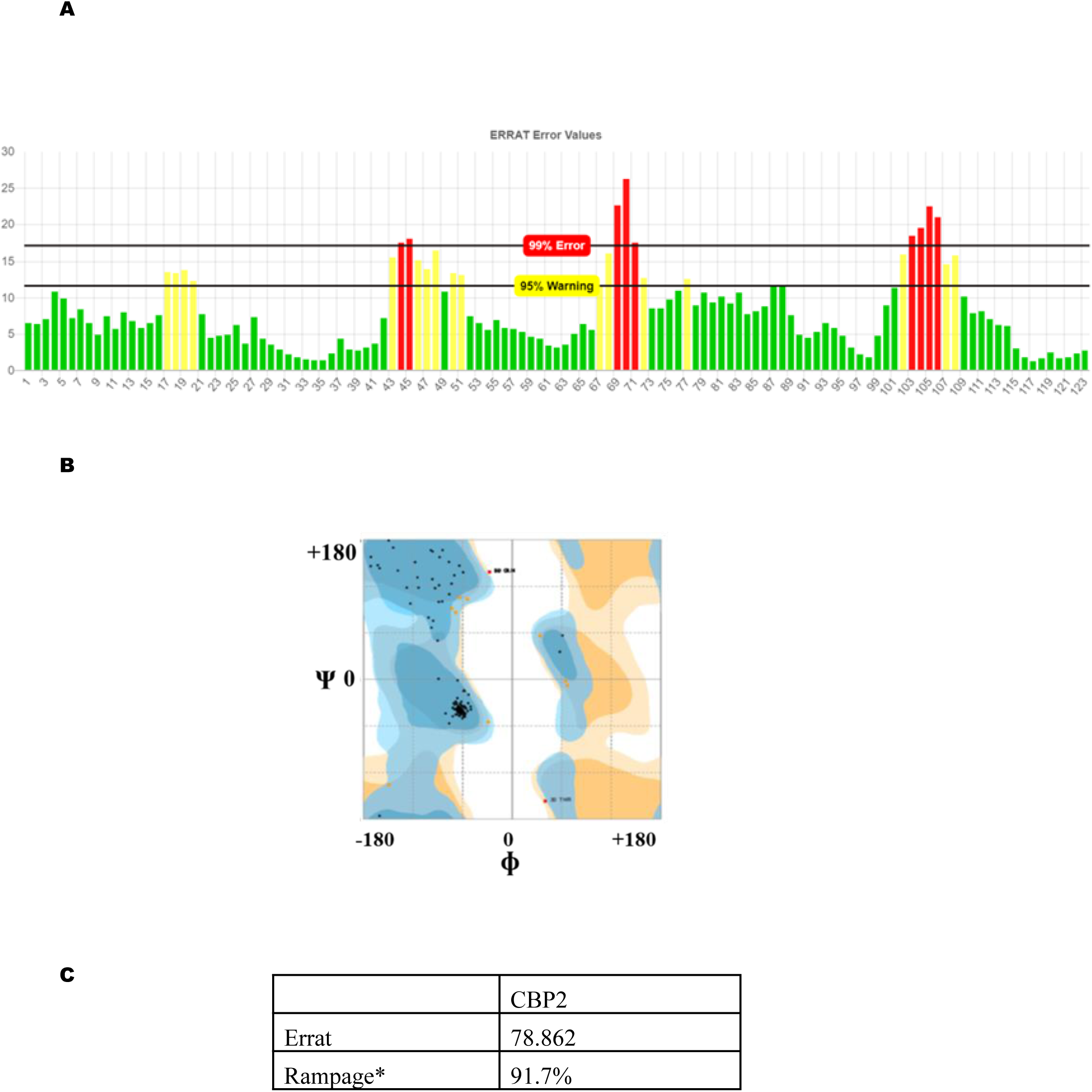
Validation of three-dimensional structure of CBP2 generated by homology modeling. (A) ERRAT (B) Ramachandran plot of generated model (C) table showing scores obtained for the theses plots.

**Fig. S3.**
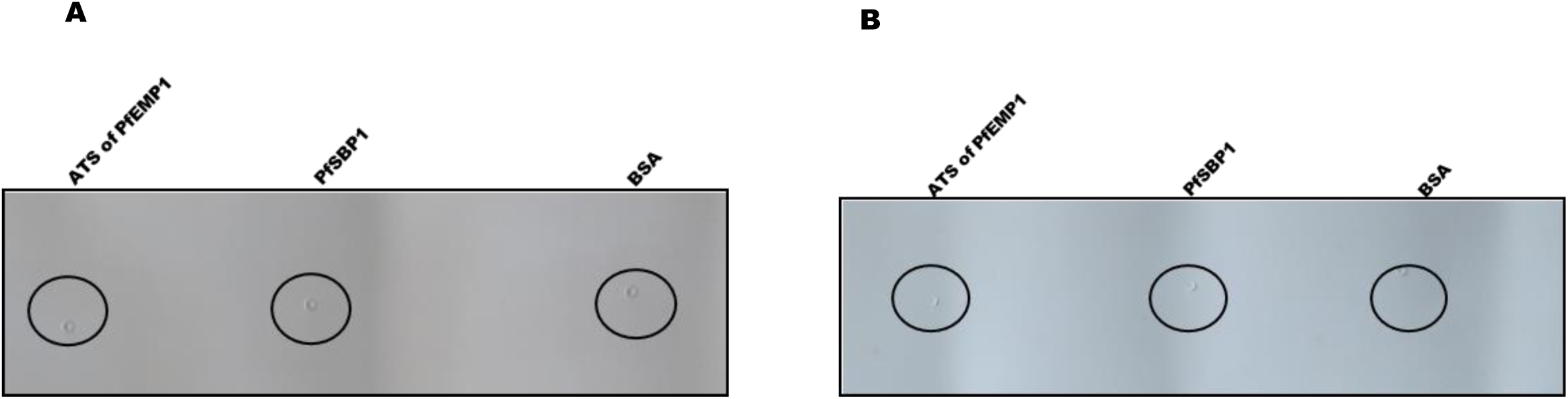
Dot-blots testing cross reactivity of anti-CBP2 antibodies and pre-immune sera with various proteins used in study. ATS of PfEMP1, PfSBP1 and BSA were spotted as labeled on the blot and probed with (A) anti-CBP2 antibodies (B) pre-immune sera of CBP2.

